# A Single Dose of Self-Transcribing and Replicating RNA Based SARS-CoV-2 Vaccine Produces Protective Adaptive Immunity In Mice

**DOI:** 10.1101/2020.09.03.280446

**Authors:** Ruklanthi de Alwis, Esther S Gan, Shiwei Chen, Yan Shan Leong, Hwee Cheng Tan, Summer L Zhang, Clement Yau, Daiki Matsuda, Elizabeth Allen, Paula Hartman, Jenny Park, Maher Alayyoubi, Hari Bhaskaran, Adrian Dukanovic, Belle Bao, Brenda Clemente, Jerel Vega, Scott Roberts, Jose A. Gonzalez, Marciano Sablad, Rodrigo Yelin, Wendy Taylor, Kiyoshi Tachikawa, Suezanne Parker, Priya Karmali, Jared Davis, Sean M. Sullivan, Steve G. Hughes, Pad Chivukula, Eng Eong Ooi

**Author notes:** Equal contribution (Ruklanthi de Alwis, Esther S Gan). Correspondence to: Dr. Eng Eong Ooi.

## Abstract

A self-transcribing and replicating RNA (STARR™) based vaccine (LUNAR^®^-COV19) has been developed to prevent SARS-CoV-2 infection. The vaccine encodes an alphavirus-based replicon and the SARS-CoV-2 full length spike glycoprotein. Translation of the replicon produces a replicase complex that amplifies and prolong SARS-CoV-2 spike glycoprotein expression. A single prime vaccination in mice led to robust antibody responses, with neutralizing antibody titers increasing up to day 60. Activation of cell mediated immunity produced a strong viral antigen specific CD8^+^ T lymphocyte response. Assaying for intracellular cytokine staining for IFN-γ and IL-4 positive CD4^+^ T helper lymphocytes as well as anti-spike glycoprotein IgG2a/IgG1 ratios supported a strong Th1 dominant immune response. Finally, single LUNAR-COV19 vaccination at both 2 μg and 10 μg doses completely protected human ACE2 transgenic mice from both mortality and even measurable infection following wild-type SARS-CoV-2 challenge. Our findings collectively suggest the potential of Lunar-COV19 as a single dose vaccine.

## INTRODUCTION

The pandemic of coronavirus disease-2019 (COVID-19) has afflicted tens of millions of people, of which hundreds of thousands have died from severe respiratory dysfunction and other complications of this disease [1]. The etiological agent of COVID-19 is the severe acute respiratory syndrome coronavirus 2 (SARS-CoV-2), which may have first emerged from a zoonotic source to then spread from person-to-person until global dissemination [1]. Current control measures to curb the pandemic, such as national lockdowns, closure of work places and schools and reduction of international travel are threatening to draw the world into a global economic recession of unprecedented scale [2]. Vaccines that elicit durable protection against SARS-CoV-2 infection are thus urgently needed [3]. Encouragingly, hundreds of different vaccine development efforts are currently in progress, some of which have even entered phase III clinical trials [4, 5].

Despite some candidates reaching late-stage clinical trials, there is some uncertainty that production can be upscaled in a sufficiently accelerated timeline to manufacture the billions of vaccine doses required to immunize the world’s population [6]. Furthermore, recent results from early phase COVID-19 vaccine trials have suggested that more than one dose would be needed to elicit reasonable levels of adaptive immune memory [7–9]. Durable protection with a single dose has been achieved with some viral live-attenuated vaccines (LAV), such as the yellow fever vaccine [10–12]. However, since the genetic determinants of the clinical fitness of SARS-CoV-2 are not well defined, development of a LAV SARS-CoV-2 strain that is safe for use in humans is challenging. An alternative approach would be to mimic the key immunogenic properties of live viral vaccines, to develop an alternate vaccine platform that could also be effective in preventing COVID-19 with a single dose. A single dose vaccine would not only avoid logistics and compliance challenges associated with multi-dose vaccines, but also allow vaccination of more individuals with each batch [6].

RNA vaccines offer a rapid approach to develop a COVID-19 vaccine [13]. RNA vaccines are designed using the genetic sequence of the viral antigen and rapidly manufactured using cell-free, rapidly scalable techniques [14]. The RNA is encapsulated in a lipid nanoparticle (LNP), which generates robust immune responses without the need for adjuvants [15, 16]. There are two main categories of RNA vaccines; 1) the conventional messenger RNA (conventional mRNA) vaccine, where the immunogen of interest is directly translated from the input vaccine transcript, and 2) the newer self-replicating RNA (replicon) vaccines [14]. Replicon vaccines encode replication machinery, usually alphavirus-based replication complex, that amplify sub-genomic RNA carrying the antigen of interest, resulting in the amplification of transcripts bearing the antigen by several orders of magnitude over the initial dose [17]. Prolonged antigen expression by such a construct could not only produce the obvious dose sparing effects [17] but potentially also elicit innate and adaptive immune responses similar to those associated with live vaccines. Herein, we show a head-to-head comparison between a self-replicating RNA vaccine using Arcturus’ proprietary Self-Transcribing and Replicating RNA (STARR™ technology and a conventional mRNA vaccine against SARS-CoV-2 and suggest that the STARR vaccine, LUNAR-COV19 offers superior vaccine-induced immune responses to conventional mRNA.

## RESULTS

### Comparison of design and expression of STARR and conventional mRNA platforms

Both LUNAR-COV19 and conventional mRNA vaccine constructs were designed to encode the full-length, unmodified, pre-fusion SARS-CoV-2 S protein (1273 aa), with LUNAR-COV19 additionally encoding the Venezuelan equine encephalitis virus (VEEV) replicase genes required for self-amplification (**Figure 1A**). We first defined the characteristics of these different constructs, which were both formulated with the same LUNAR LNP lipid formulation. Despite differences in RNA lengths for LUNAR-COV19 and conventional mRNA, the LNP diameter, polydispersity index and RNA trapping efficiency were similar (**Figure 1B**). *In vitro* expression of the LUNAR-COV19 and conventional mRNA vaccine were confirmed in cell lysate 24 hours post-transfection through positive western blot detection of the S protein (**Figure 1C**). It was also observed that both vaccines expressed a mixture of full-length S protein and cleaved S protein, i.e. into S1 and S2 transmembrane and cytoplasmic membrane domains (**Figure 1C**). We then compared *in vivo* protein expression of the two RNA platforms in BALB/c mice, by using STARR and conventional mRNA constructs that expressed a luciferase reporter (**Figure 1D**). As expected, animals injected with the conventional mRNA vaccine construct showed high *in vivo* luciferase expression at day 1 although the expression levels declined significantly three days post injection. In contrast, the luciferase expression in STARR injected mice showed increased signal of protein production compared to conventional mRNA at all time points after Day 1 up to Day 7 post-inoculation (the last time point measured) and at doses ≥2.0 μg, protein expression appeared to be still rising at day 7 (**Figure 1D**). These data showed that dose-for-dose, the STARR luciferase construct yielded higher and more prolonged duration of luciferase expression compared to mice injected with the conventional mRNA luciferase construct.

**Figure 1.**
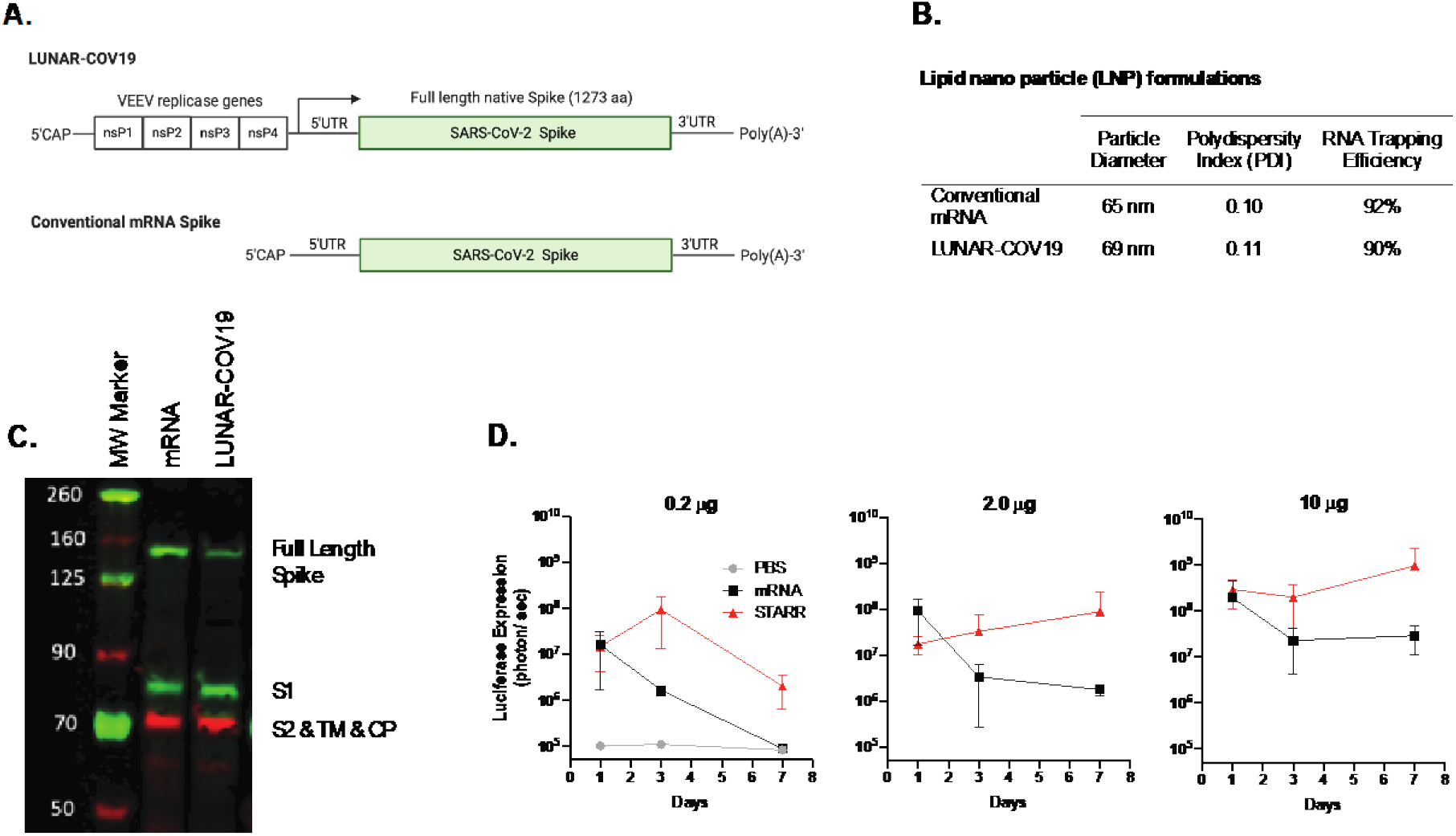
Design and Expression of a SARS-COV-2 vaccine with conventional mRNA and self-transcribing and replicating RNA (STARR^®^) platforms. **A)** Schematic diagram of the SARS-CoV-2 self-replicating STARR RNA (LUNAR^®^-COV19) and conventional mRNA vaccine constructs. The STARR construct encodes for the four non-structural proteins, ns1-ns4, from Venezuelan equine encephalitis virus (VEEV) and the unmodified full-length pre-fusion spike (S) protein of SARS-CoV-2. The mRNA construct also codes for the same SARS-CoV-2 full length spike S protein. **B)** Physical characteristics and RNA trapping efficiency of the LNP encapsulating conventional mRNA and LUNAR-COV19 vaccines. **C)** Western blot detection of SARS-CoV-2 S protein following transfection of HEK293 cells with LUNAR-COV19 and conventional mRNA. **D)** *In vivo* comparison of protein expression following IM administration of LNP containing luciferase-expressing STARR RNA or conventional mRNA. Balb/c mice (*n*=3/group) were injected IM with 0.2 μg, 2.0 μg and 10.0 μg of STARR RNA or conventional mRNA formulated with the same lipid nanoparticle. Luciferase expression was measured by *in vivo* bioluminescence on days 1, 3 and 7 post-IM administration. S domain 1 = S1, S domain 2 = S2, transmembrane domain = TM, cytoplasmic domain = CP; aka = also known as.

### Immune gene expression following LUNAR-COV19 and conventional mRNA vaccination

C57BL/6J mice were vaccinated with LUNAR-COV19 or conventional mRNA vaccines at 0.2 μg, 2 μg and 10 μg doses or PBS control. No significant mean loss in animal weight occurred over the first 4 days, except for those that received 10 μg of LUNAR-COV19 (**Figure 2A)**. However, apart from weight loss, there were few other clinical signs as indicated by the minimal differences in clinical scores. Both weight and clinical scores improved uneventfully after day 3 post vaccination.

**Figure 2:**
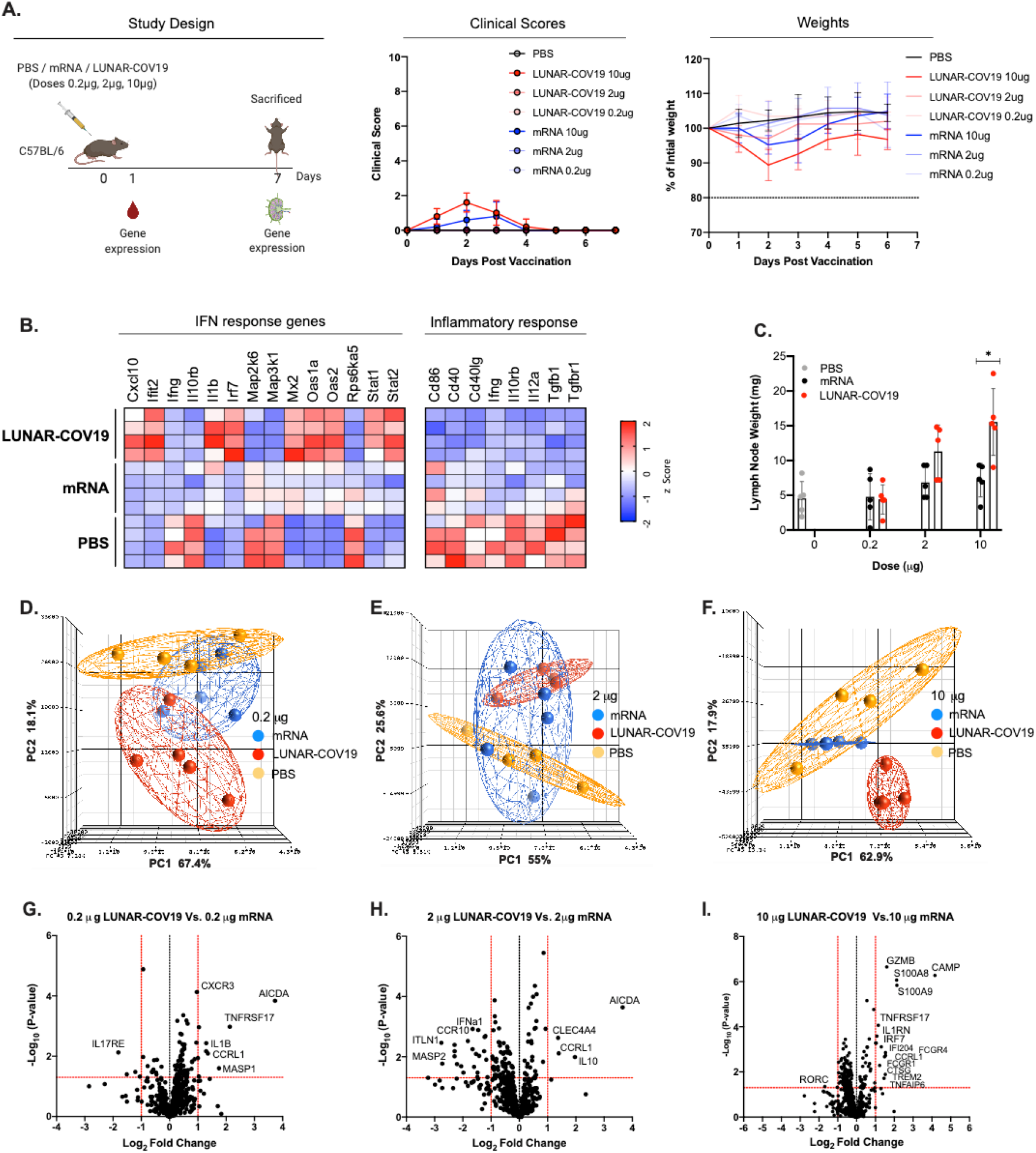
Clinical Scores, mouse weights and transcriptomic analysis of immune genes following vaccination with LUNAR-COV19 or conventional mRNA SARS-CoV-2 vaccine candidates. **A)** C57BL/6 mice (*n*=5/group) were immunized with either PBS, mRNA or LUNAR-COV19 (doses 0.2 μg, 2 μg or 10 μg), weight and clinical scores assessed every day, bled at day 1 post-immunization, sacrificed at 7 days post-vaccination and lymph nodes harvested. Gene expression of inflammatory genes and immune genes were measured in whole blood (at day 1) and lymph nodes (at day 7), respectively. **B**) Expression of IFN and inflammatory response genes in whole blood presented as heatmap of z scores. **C)** Lymph node weights at 7 days post-vaccination. Principal component analysis (PCA) of immune gene expression following vaccination with conventional mRNA or LUNAR-COV19 at doses **D)** 0.2 μg, **E)** 2 μg and **F)** 10 μg. Volcano plots of fold change of LUNAR-COV19 versus conventional mRNA (x-axis) and Log_10_ *P*-value of LUNAR-COV19 versus conventional mRNA (y-axis) for doses **G)** 0.2 μg, **H)** 2 μg and **I)** 10 μg. Study design schematic diagram created with BioRender.com. Weights of lymph nodes were compared between groups using a two-tailed Mann-Whitney *U* test with * denoting 0.05<*P*<0.01.

The innate immune response, particularly the type-I interferon (IFN) response has previously been shown to be associated with vaccine immunogenicity following yellow fever vaccination [11, 12, 18]. Furthermore, we have also found that reactive oxygen species-driven pro-inflammatory responses underpinned systemic adverse events in yellow fever vaccination [19, 20]. Therefore, we measured the expression of innate immune and pro-inflammatory genes in whole blood of C57BL/6 mice inoculated with either PBS, conventional mRNA vaccine or LUNAR-COV19. Genes in the type-I IFN pathway were the most highly expressed in animals inoculated with LUNAR-COV19 compared to either conventional mRNA vaccine or PBS (**Figure 2B** and **Supplementary Figure 1**). By contrast, genes associated with pro-inflammatory responses were mostly reduced in abundance following LUNAR-COV19 vaccination compared with either conventional mRNA vaccine or PBS (**Figure 2B** and **Supplementary Figure 1**).

Since adaptive immune responses develop in germinal centers in the draining lymph nodes, we dissected the draining lymph nodes at day 7 post-inoculation (study schematic in **Figure 2A**). The inguinal lymph nodes of mice inoculated with LUNAR-COV19 showed a dose-dependent increase in weight, unlike those from mice inoculated with either conventional mRNA vaccine or PBS; the mean weight of lymph nodes from mice given 10 μg of LUNAR-COV19 was significantly higher than those given the equivalent conventional mRNA vaccine (**Figure 2C**). Principal component analysis (PCA) of immune gene expression showed clustering of responses to each of the 3 doses of LUNAR-COV19 away from the PBS control (depicted as red and orange spheres in **Figure 2D-F**), indicating clear differences in immune gene expression between LUNAR-COV19 vaccinated and placebo groups. These trends were also dissimilar to those from mice given conventional mRNA vaccine where at all tested doses, the PCA displayed substantial overlap with PBS control (shown as blue and orange spheres in **Figure 2D-F**).

We next assessed the differentially expressed genes in the lymph nodes of mice given LUNAR-COV19 compared to those inoculated with mRNA vaccine. Volcano plot analysis identified significant upregulation of several innate, B and T cells genes in LUNAR-COV19 immunized animals (**Figure 2G-I**). Some of the most highly differentially expressed genes included, GZMB (required for target cell killing by cytotoxic immune cells) [21], S100A8 and S100A9 (factors that regulate immune responses via TLR4) [22], TNFRSF17 (also known as BCMA and regulates humoral immunity) [23], CXCR3 (chemokine receptor involved in T cell trafficking and function) [24] and AICDA (mediates antibody class switching and somatic hypermutation in B cells) [25]. These findings collectively indicate that the adaptive immune responses in the draining lymph nodes of mice inoculated with LUNAR-COV19 may differ to those given the non-replicating mRNA vaccine.

### LUNAR-COV19 induced robust T cell responses

We next investigated the cellular immune response following vaccination of C57BL/6 mice (*n*=5 per group) with LUNAR-COV19 or conventional mRNA. At day 7 post-vaccination, spleens were harvested and assessed for CD8 and CD4 T cells by flow-cytometry. The CD8+ T cell CD44+CD62L-effector/memory subset was significantly expanded in LUNAR-COV19 vaccinated mice compared to those given either PBS or conventional mRNA vaccine (**Figure 3A-B**). There was no statistically significant difference in the proportion of CD4+ T effector cells in these animals (**Figure 3C**). IFNγ+ CD8+ T cells (with 2 μg and 10 μg doses) and IFNγ+ CD4+ T cells (in 0.2 μg and 10 μg) were proportionately higher, as found using intracellular staining (ICS) with flow cytometry, in LUNAR-COV19 as compared to conventional mRNA vaccinated animals (**Figure 3D-F**).

**Figure 3.**
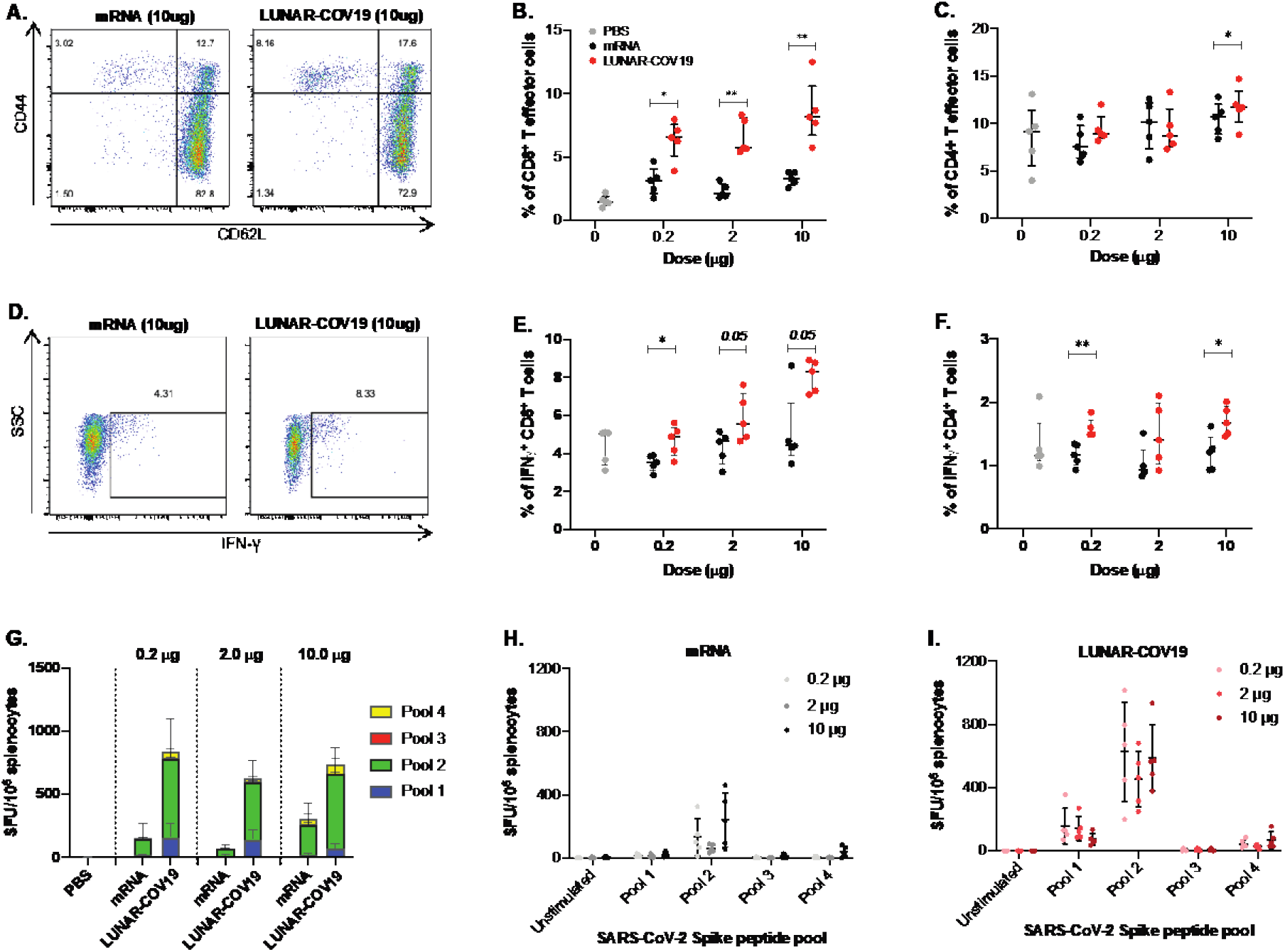
Cellular immune responses following vaccination with LUNAR-COV19 and conventional mRNA. C57BL/6 mice (*n*=5 per group) were immunized with 0.2 μg, 2.0 μg, or 10.0 μg of LUNAR-COV19 or conventional mRNA via IM, sacrificed at day 7 post-vaccination and spleens analyzed for cellular T cell responses by flow-cytometry and ELISPOT. **A-B**) CD8^+^ and **C**) CD4^+^ T effector cells were assessed in vaccinated animals using surface staining for T cell markers and flow-cytometry. **D-E**) IFNγ^+^ CD8^+^ T cells and **F**) Ratio of IFNγ^+^/ IL4^+^ CD4^+^ T cells in spleens of immunized mice were assessed following *ex vivo* stimulation with PMA/IO and intracellular staining. **G-I**) SARS-CoV-2 S protein-specific responses to pooled S protein peptides were assessed using IFNγ ELISPOT assays following vaccination with mRNA (**H**) or LUNAR-COV19 (**I**). Percentage of CD8+ cells, CD4+ cells, IFNγ and IL4 producing T cells were compared between groups using two-tailed Mann-Whitney *U* test with * denoting 0.05<*P*<0.01, and **0.01<*P*<0.001.

SARS-CoV-2 specific cellular responses were assessed in vaccinated animals by ELISPOT. A set of 15-mer peptides covering the full length SARS-CoV-2 S protein were divided into 4 pools and tested for IFNγ+ responses in splenocytes of vaccinated and non-vaccinated animals. SARS-CoV-2-specific cellular responses (displayed as IFNγ+ SFU/10^6^ cells) were detected by ELISPOT in both LUNAR-COV19 and conventional mRNA vaccine immunized animals compared to PBS control (**Figure 3G-I**). These responses were substantially higher across all doses in LUNAR-COV19 compared to conventional mRNA vaccinated groups (**Figure 3G-I**). Even the highest tested dose (10 μg) of conventional mRNA vaccine produced IFNγ+ ELISPOT responses that were appreciably lower than those by the lowest dose (0.2 μg) of LUNAR-COV19.

### LUNAR-COV19 induced superior humoral immune responses

SARS-CoV-2-specific humoral responses following vaccination with a single injection were characterized in two different mouse models, BALB/c and C57BL/6. Female mice (*n*=5 per group) were vaccinated at day 0 and bled every 10 days, up to day 60 for BALB/c and day 30 for C57BL/6 (**Figure 4A**). SARS-CoV-2 S-specific IgM responses were tested at 1:2000 serum dilution using an in-house Luminex immuno-assay. All tested doses of the conventional mRNA vaccine and LUNAR-COV19 produced detectable S-specific IgM responses in both mouse models (**Figure 4B-C**). When comparing LUNAR-COV19 to conventional mRNA vaccinated BALB/c mice, no difference in IgM responses was observed; IgM levels in C57BL/6 mice were higher in LUNAR-COV19 vaccinated C57BL/6 mice at day 10 post vaccination. In contrast, SARS-CoV-2 S-specific IgG (at 1:2000 serum dilution) levels were universally higher from day 20 onwards in animals inoculated with LUNAR-COV19 compared to conventional mRNA vaccine (**Figure 4D-E**). Perhaps even more remarkably, in BALB/c vaccinated with LUNAR-COV19, the IgG levels continued to increase until day 50 post-vaccination; C57BL/6 mice were only monitored until day 30 post-vaccination. This trend contrasted sharply with mice that received the conventional mRNA vaccine where in BALB/c mice antibody levels plateaued after day 10 post-vaccination; although increasing S-specific IgG levels were observed in conventional mRNA-vaccinated C57BL/6 mice these were universally lower than those that received LUNAR-COV19.

**Figure 4:**
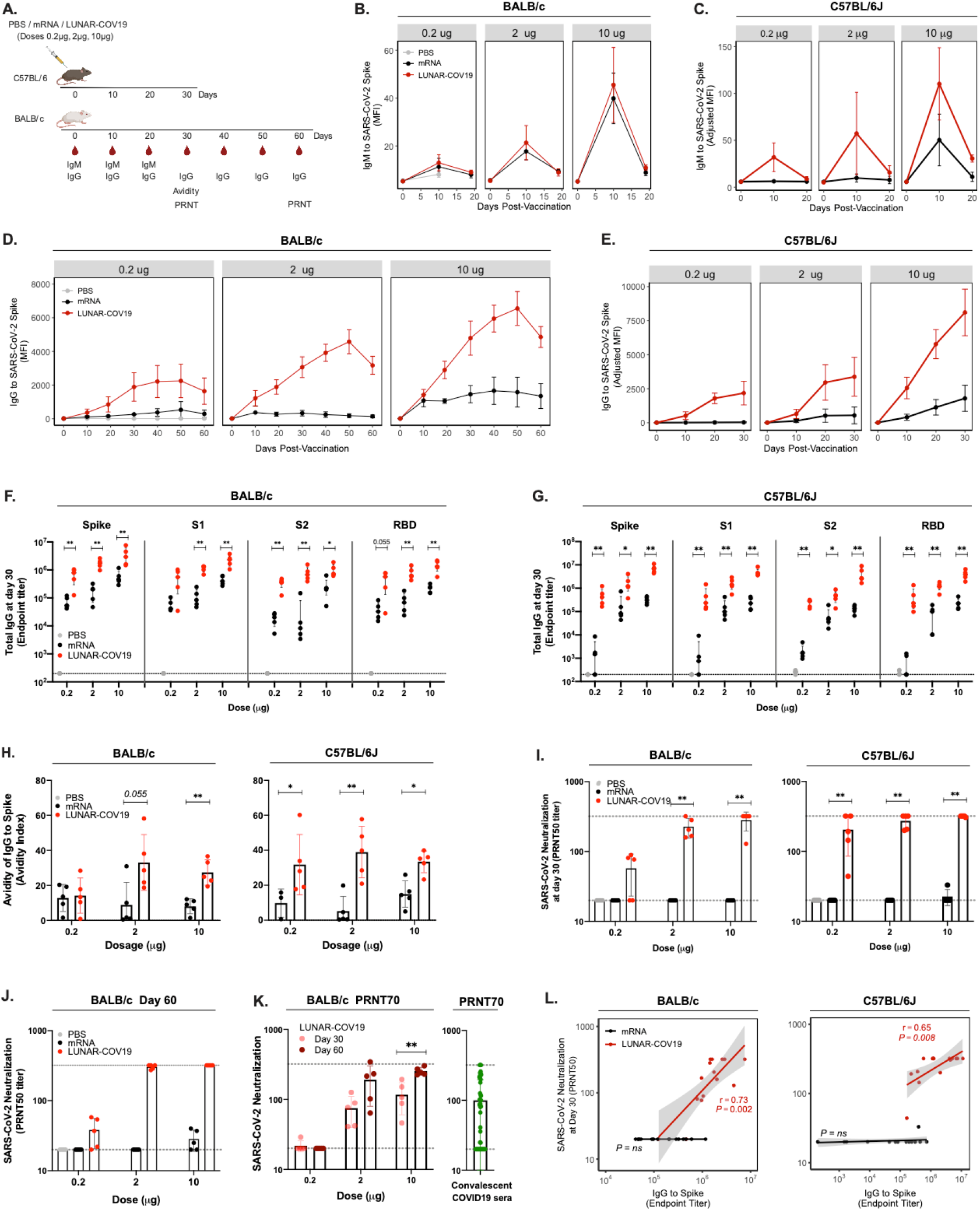
LUNAR-COV19 elicits a higher quality humoral response than conventional mRNA platform. **A)** BALB/c and C57BL/6J mice were immunized via IM with 0.2 μg, 2μg, or 10 μg of LUNAR-COV19 or conventional mRNA (*n*=5/group). Blood sampling was conducted at baseline, and days 10, 19, 30, 40, 50 and 60 post-vaccination for BALB/c and days 10, 20 and 30 for C57BL/6J. **B**-**C**) IgM and **D**-**E**) IgG against the SARS-CoV-2 S protein over time, assessed using insect cell-derived whole S protein in a Luminex immuno-assay (measured as MFI). IgG endpoint titers to mammalian-derived whole S protein, S1, S2 and RBD proteins to mammalian-derived whole S protein at day 30 post-vaccination were assessed in **F)** BALB/c and **G)** C57BL/6J. **H**) Avidity of SARS-CoV-2 S protein-specific IgG at day 30 post-immunization was measured using 8M urea washes. **I**) Neutralizing antibody (PRNT_50_ titers) at day 30 post-vaccination against a clinically isolated live SARS-CoV-2 virus measured in both BALB/c and C57BL/6J. Gray dashed lines depict the serum dilution range (i.e. from 1:20 to 1:320) tested by PRNT. **J**) PRNT50 and **K**) PRNT70 of SARS-CoV-2 neutralization at day 30 and day 60 post-vaccination in BALB/c and convalescent sera from COVID-19 patients. **L**) Correlation analysis of Spike-specific IgG endpoint titers against SARS-CoV-2 neutralization (PRNT50). Antibody data were compared between groups using a two-tailed Mann-Whitney *U* test with * denoting 0.05<*P*<0.01, and **0.01<*P*<0.001.

In depth characterization of the SARS-CoV-2 specific IgG response in vaccinated animals was conducted at day 30 post-immunization to assess which regions of S protein are targeted. IgG endpoint titers were estimated to full ectodomain S protein, S1, S2 and receptor binding domain (RBD) regions. As expected for both vaccine candidates the majority of SARS-CoV-2 specific IgG recognized S1, which contains the RBC, although high IgG endpoint titers were also detected to S2 protein (**Figure 4F-G**). However, LUNAR-COV19 elicited IgG endpoint titers were universally and significantly higher compared to those produced by conventional mRNA vaccination (**Figure 4F-G**). Notably, IgG that bind the RBD of S protein, which is an immunodominant site of neutralizing antibodies [26, 27], were also higher in LUNAR-COV19 compared to conventional mRNA vaccinated animals. It is also noteworthy that at lower doses, conventional mRNA vaccine but not LUNAR-COV19 struggled to elicit high SARS-CoV-2 specific IgG titers in the more Th1 dominant C57BL/6 mouse strain (**Figure 4G**). Taken collectively, a single dose of LUNAR-COV19 induced significant differences in immune gene expression and superior cellular immune responses in draining lymph nodes compared to the conventional mRNA vaccine and consequently greater and more prolonged humoral immune responses.

We assessed both the binding strength (avidity) and the neutralizing ability of the antibody response elicited by these vaccine constructs. Serum IgG avidity was measured at day 30 post-vaccination using a modified Luminex immuno-assay with 8M urea washes. LUNAR-COV19 elicited higher avidity S protein-specific IgG in both mouse models at all tested doses (**Figure 4H**). These differences were observed, with the exception of 0.2 μg in BALB/c, across all doses (**Figure 4H**), indicating that LUNAR-COV19 elicited better quality antibodies, suggesting superior affinity maturation with the LUNAR-COV19 vaccine.

Neutralization of live SARS-CoV-2 by serum from vaccinated animals was assessed using the plaque reduction neutralization test (PRNT). At day 30 LUNAR-COV19 vaccinated BALB/c mice showed a clear dose-dependent elevation in PRNT_50_ titers; 4 out of 5 (80%) of mice in the 10 μg LUNAR-COV19 group showed PRNT_50_ titers greater than 320, which was the upper limit of our dilution (**Figure 4I**). Similar dose-dependent trends in PRNT_50_ titers were also found in C57BL/6 mice although in these animals, the PRNT_50_ titers of several animals exceeded 320 even with the lowest 0.2 μg dose vaccination (**Figure 4I**). In sharp contrast, PRNT_50_ titers in animals inoculated with the conventional mRNA vaccine construct were, except for one C57BL/6J mouse that received 10 μg dose, all <20 (**Figure 4I**). Unexpectedly but encouragingly, PRNT_50_ and PRNT_70_ titers of LUNAR-COV19 vaccinated BALB/c mice continued to rise between day 30 and day 60 after a single vaccination (**Figure 4J-K**) and at both time points for doses ≥2.0 μg. These titers were comparable to PRNT_70_ titers for sera from convalescent COVID-19 patients (**Figure 4K**).

We also found that the S protein IgG titers positively correlated with PRNT_50_ titers with LUNAR-COV19 vaccinated mice in both mouse models (**Figure 4L**). Similar positive correlations were also observed with IgG against S1 and RBD (**Supplementary Figure 1**). By contrast, we found no correlation between IgG and PRNT_50_ titers in conventional mRNA vaccinated mice (**Figure 4L**). Taken collectively, our antibody response analyses suggest that the higher PRNT_50_ titers following vaccination with LUNAR-COV19 are not only strongly associated with the amount of IgG produced but are also a factor of the superior quality of the anti-SARS-CoV-2 antibodies produced following vaccination with LUNAR-COV19.

### LUNAR-COV19 vaccination showed a Th1 dominant response

A safety concern for a coronavirus vaccine is the risk of vaccine-associated immune enhancement of respiratory disease (VAERD) [28]. Indeed, SARS-CoV and MERS-CoV vaccine development have highlighted the importance of Th1 skewed responses in mitigating the risk of vaccine-induced immune enhancement [29, 30]. Therefore, we investigated the Th1/ Th2 balance elicited by vaccination with both conventional mRNA and LUNAR-COV19. The IgG subclass fate of plasma cells are highly influenced by T helper (Th) cells [31]. At day 30 post-vaccination, both conventional mRNA and LUNAR-COV19, induced comparable amounts of SARS-CoV-2 S-specific IgG1, a Th2-associated IgG subclass in mice, except for the 0.2 μg dose in C56BL/6J mice (**Figure 5A-B**). In contrast, the Th1-associated IgG subclasses - IgG2a in BALB/c and IgG2c in C56BL/6J - were significantly greater in LUNAR-COV19 vaccinated animals. The ratios of S protein-specific IgG2a/IgG1 (Balb/c) and IgG2c/IgG1 (C57BL/6) were greater than 1 in LUNAR-COV19 vaccinated animals (**Figure 5A-B)**. Except for the 0.2 μg dose, these ratios were all significantly greater with LUNAR-COV19 compared to the conventional mRNA vaccinated animals.

**Figure 5.**
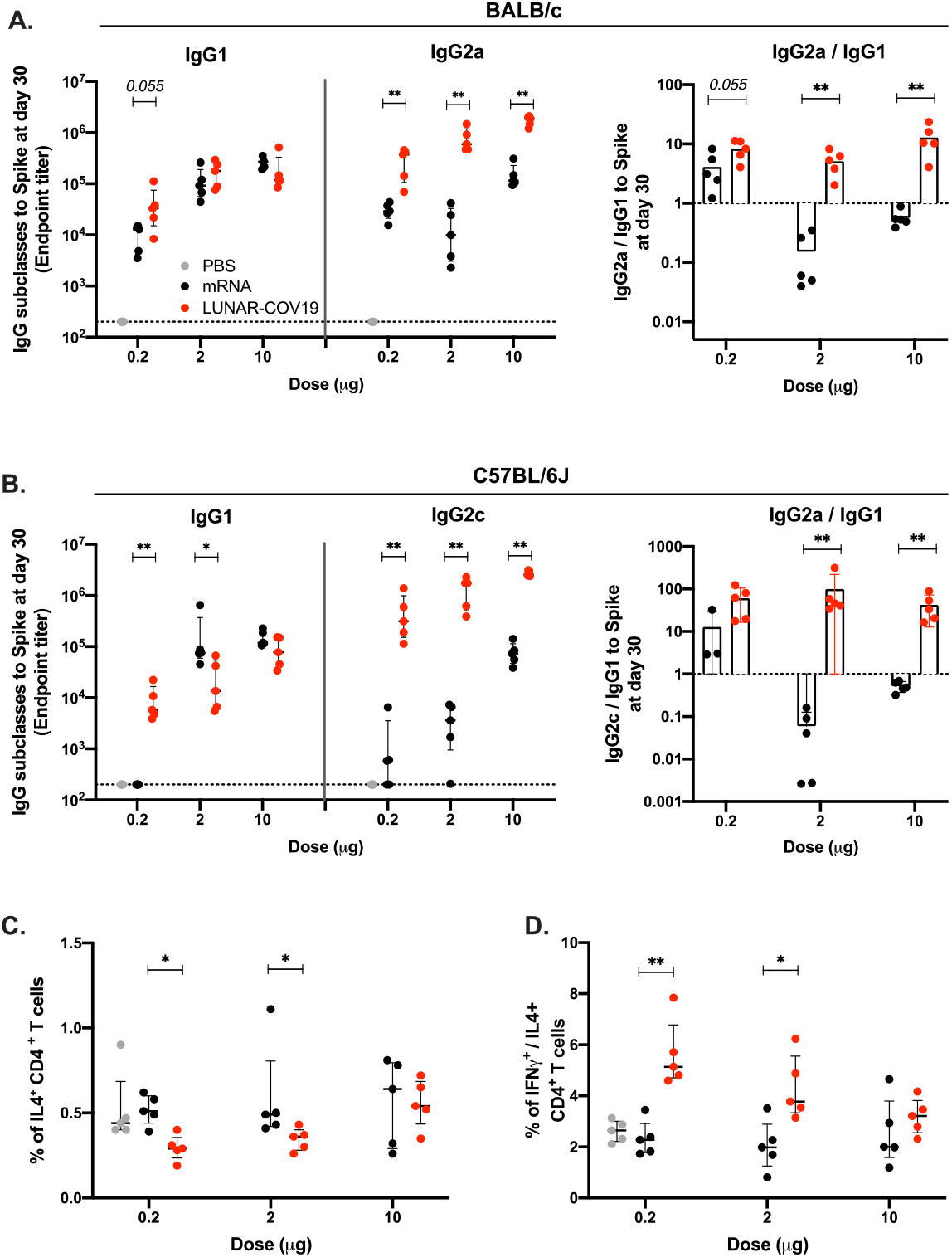
LUNAR-COV19 elicits Th1 biased immune responses. SARS-CoV-2 spike-specific IgG subclasses and the ratio of IgG2a/c/IgG1 at 30 days post-vaccination with LUNAR-COV19 and conventional mRNA in **A**) BALB/c and **B**) C57BL/6J mice. Th2 cytokine and Th1/Th2 skew in CD4 T cells at day 7 post-vaccination in C57BL/6J mice measured by ICS as **C**) percentage of IL4+ CD4 T cells and **D**) ratio of IFNγ^+^/IL4^+^ CD4^+^ T cells. Antibody titers and T cell data were compared between groups using a two-tailed Mann-Whitney *U* test with * denoting 0.05<*P*<0.01, and **0.01<*P*<0.001.

Additionally, we used ICS to investigate the production of IFNγ (Th1 cytokine) and IL4 (Th2 cytokine) by CD4+ T cells in spleens at day 7 post vaccination C56BL/6J mice. As was described above, compared to conventional mRNA vaccination, IFNγ levels were significantly greater in LUNAR-COV19 vaccinated animals (Figure 3F). IL4 expression in CD4 T cells were slightly higher with conventional mRNA as compared to LUNAR-COV19 at 0.2 and 2.0 μg doses (**Figure 5C**). In comparing the IFNγ and IL4 levels in individual mice, we found that the ratios of IFNγ/IL4 in CD4+ T cells for both LUNAR-COV19 and conventional mRNA vaccinated mice were universally above 1 (**Figure 5D**). The ratio of IFNγ/IL4 in CD4+ T cells in mice given the 0.2 and 2.0 μg doses were significantly greater with LUNAR-COV19 than conventional mRNA vaccination (**Figure 5F**). However, the elevated ratios at these doses were due to a decrease in IL4 expression at levels below background (i.e. PBS control mice), rather than reduced IFNγ and hence Th1 activity. Taken collectively, our data show that LUNAR-COV19 produced a Th1 biased adaptive immune response.

### Single dose of LUNAR-COV19 protects from a lethal infection of SARS-CoV-2

Finally, we tested the efficacy of LUNAR-COV19 in protecting against infection and mortality in a lethal SARS-CoV-2 challenge model. Transgenic hACE2 mice immunized with either PBS, or 2 μg or 10 μg of LUNAR-COV19 vaccine were intranasally challenged with live SARS-CoV-2 virus (5×10^4^TCID_50_) at day 30 post-vaccination. This was the same isolate as that used for our PRNT assays. Mice were then divided into two groups: one group was tracked for weight, clinical scores and survival; a second group of mice were euthanized at 5 days post injection (dpi) and viral loads assessed in the respiratory tract (trachea to lung) and brain (**Figure 6A**). Measurement of PRNT_70_ titers confirmed the generation of neutralizing antibodies in LUNAR-COV19-vaccinated hACE2 mice (**Figure 6B**). Irrespective of tested dosages, mice that received the LUNAR-COV19 vaccine showed unchanged weight and no clinical sign, while the PBS mice showed significant drop in weight and increased clinical scores upon challenge with wild-type SARS-CoV-2 (**Figure 6C-D**). LUNAR-COV19 vaccination at both 2 μg and 10 μg doses fully protected hACE2 mice from an otherwise 100% mortality at day 7 post-challenge (**Figure 6E**). Assessment of tissue viral load at day 5 post-challenge found minimal to no SARS-CoV-2 RNA (**Figure 6F**) in contrast to unvaccinated animal controls. Although viral RNA was detectable at very low levels in some animals, this was not associated with any presence of infectious viral particles, so most like represents viral RNA fragments rather than intact viral RNA genomes. No detectable infectious virus was found in either the respiratory tracts or brains of LUNAR-COV19 vaccinated animals (**Figure 6G**). By contrast, unvaccinated animals showed 4 and 8 logs of infectious SARS-CoV-2 in the respiratory tract and brain, respectively (**Figure 6G**). Collectively, these data show that a single dose of LUNAR-COV19 vaccine induced robust humoral and cellular immune responses that led to complete protection of hACE2 mice from a lethal SARS-CoV-2 challenge.

**Figure 6.**
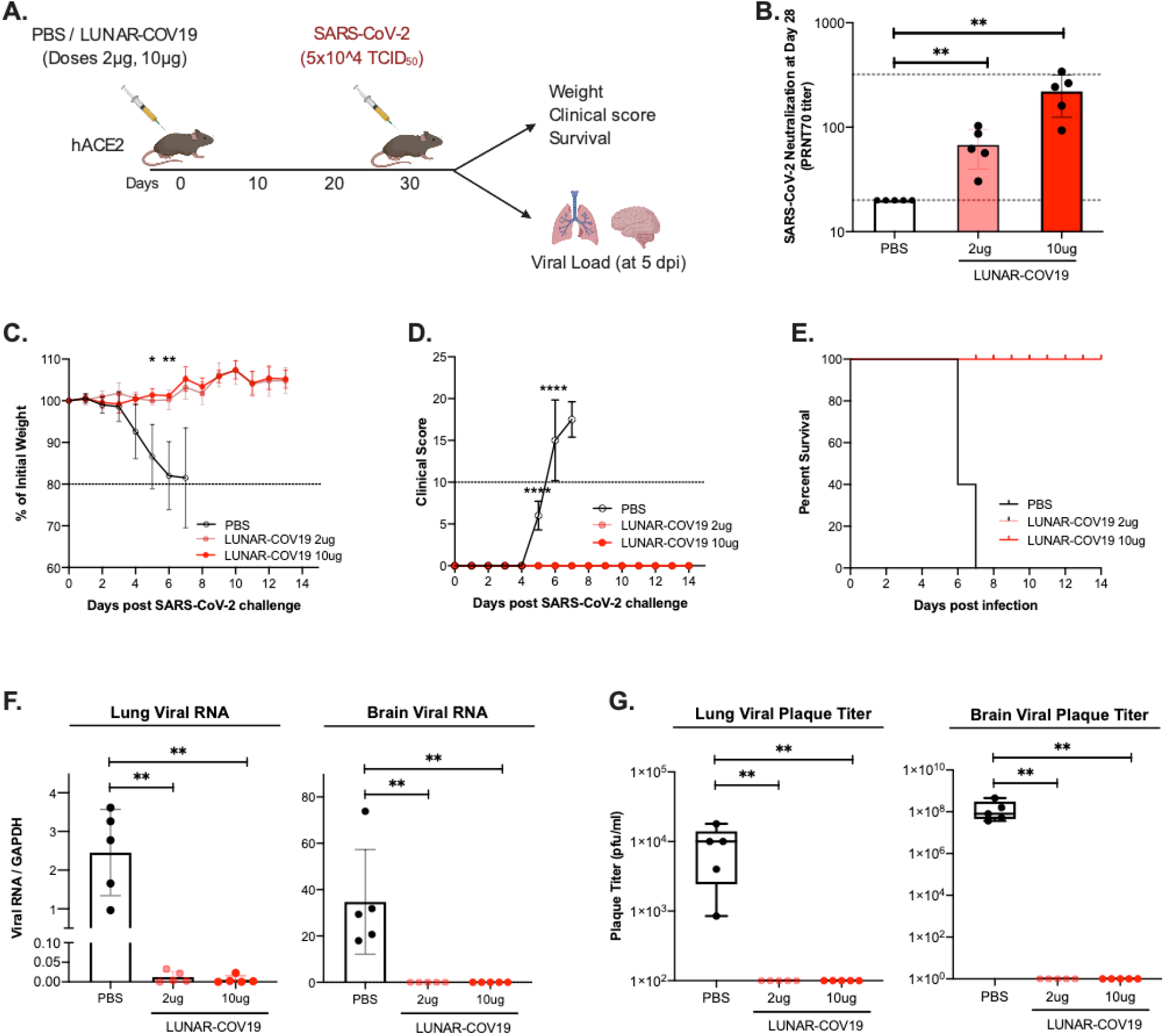
Single dose of LUNAR-COV19 protects hACE2 mice against a lethal challenge of SARS-CoV-2 virus. **A**) hACE2 transgenic mice were immunized with a single dose of either PBS or 2 μg or 10 μg of LUNAR-COV19 (*n*=5 per group), then challenged with live SARS-CoV-2 at 30 days post-vaccination, and assessed for either survival (with daily weights and clinical scores) or sacrificed at day 5 post-challenge and measured lung and brain tissue viral loads. Study design schematic diagram was created with BioRender.com **B**) Live SARS-CoV-2 neutralizing antibody titers (PRNT70) measured at 28 days post-vaccination. **C**) Weight, **D**) clinical score and **E**) survival was estimated following challenge with a lethal dose (5×10^5 TCID_50_) of live SARS-CoV-2 virus. **F**) Viral RNA and **G**) infectious virus in the lungs and brain of challenged mice were measured with qRT-PCR or plaque assay, respectively. PRNT_70_ and viral titers (RNA and plaque titers) were compared across groups using the non-parametric Mann-Whitney *U* test. Weights and clinical scores at different timepoints were compared between PBS and 10ug LUNAR-CoV19 immunized mice using multiple *t*-tests. *P*-values are denoted by * for 0.05<*P*<0.01, ** for 0.01<*P*<0.001, *** for 0.001<*P*<0.0001, **** *P*<0.00001.

## DISCUSSION

The pandemic of COVID-19 has necessitated rapid development of vaccines. Encouragingly, several COVID-19 vaccine candidates are now in clinical trials and more are entering first-in-human trials. However, the majority of vaccine candidates being developed require two or more doses for sufficient adaptive immune responses. Requirement for a second shot could complicate compliance rate in mass vaccination campaigns and results in fewer subjects vaccinated per batch, thereby reducing the efficiency of vaccination. Hence, a single dose vaccine that generates robust and sustained cellular and humoral immunity, without elevating the risk of vaccine-mediated immune enhancement, remains an unmet need.

Amongst the licensed vaccines for other diseases, live attenuated vaccines can offer the most durable protection against viral diseases. Live vaccines infect and replicate at sites of inoculation and some even in draining lymph nodes. Replication enables endogenous and sustained expression of viral antigens that enable antigen presentation to stimulate cytotoxic CD8+ T cells. Expressed antigens taken up by antigen presenting cells also trigger CD4+ T cell help that drives affinity maturation in B cells. Studies on the live attenuated yellow fever vaccine, have shown that a longer period of stimulation of the adaptive immune response results in superior adaptive immune responses [32]. Although work to determine which of these correlates of live vaccines are mechanistic determinants of adaptive immunity is still ongoing, the ability of self-replicating RNA vaccines to simulate the sustained antigen presentation characteristics of live vaccination could offer durable immunity against COVID-19.

Numerous studies have shown RNA vaccines to be immunogenic. In this study, we conducted a side-by-side comparison of the immunogenicity elicited by two SARS-CoV-2 RNA vaccine candidates, a conventional mRNA construct and the STARR construct, LUNAR-COV19. We found that, compared to conventional mRNA, LUNAR-COV19 produced higher and longer protein expression *in vivo*, upregulated the gene expression of several innate, B and T cell response genes in the blood and draining lymph nodes. These properties were associated with significantly greater neutralizing antibody and SARS-CoV-2 specific IgG responses, CD8+ T cell responses, IFNγ+ ELISPOT responses, and Th1 skewed responses (which have been shown to associate with protection from VAERD) than conventional mRNA. Interestingly, despite the highest tested dose of conventional mRNA eliciting comparable S protein-specific antibodies as the lowest tested dose of LUNAR-COV19, the conventional mRNA-elicited IgG did not show such robust avidity or neutralization activity as those from LUNAR-COV19 vaccination. These data suggest a qualitatively better humoral immune response with superior affinity maturation of B-cells with the LUNAR-COV19 vaccine. Our findings thus highlight the immunological advantages of self-replicating RNA over conventional mRNA platforms.

The superior quality of immune responses elicited by LUNAR-COV19 over the conventional mRNA vaccine construct could be attributable to multiple factors. Higher and longer expression of immunogens produce better immunity [32], likely through better engagement of T follicular helper cells and thereby leading to more diverse antibody targets and more robust neutralizing antibody responses [33, 34]. Replication of LUNAR-COV19 results in the formation of a negative-strand template for production of more positive-strand mRNA and sub-genomic mRNA expressing the S transgene. Interaction between the negative- and positive-strands forms a double stranded RNA (dsRNA) intermediate, which would interact with TLR3 and RIG-I-like receptors to stimulate type 1 interferon responses [35–37], which we and others have previously shown to correlate with superior adaptive immune responses [11, 12, 18]. Production of IFNγ can also stimulate development of cytotoxic CD8+ T cells [36]. Importantly, the S protein does contain human CD8+ T cell epitopes. As suggested by recent findings on T cell responses to SARS-CoV-2 and other coronavirus infections [38–40], the development of T cell memory could be important for long-term immunity.

It is unclear whether the VEEV nsP1-4 forming the replication complex contains any immunogenic properties although mutations in the nsP proteins have been shown to affect the induction of type I IFN [41]. Although unexplored in our current study, VEEV replicons have also been shown to adjuvant immune responses at mucosal sites [42], further justifying the use of STARR platform to develop a COVID-19 vaccine.

In conclusion, STARR vaccine platform as exemplified by LUNAR-COV19, offers an approach to simulate key immunogenic properties of live virus vaccination and offers the potential for an effective single-shot vaccination against COVID-19.

## METHODS (Supplement 1)

### Vaccine plasmid constructs and design

A human codon-optimized spike (S) glycoprotein gene of SARS-CoV-2 (GenBank accession: YP_009724390) was cloned into plasmids pARM2922 and pARM2379 for generation of SARS-CoV-2 Spike expressing STARR and conventional mRNA, respectively. The STARR plasmid also encoded for the Venezuela equine encephalitis virus (VEEV) non-structural proteins nsP1, nsP2, nsP3 and nsp4, which together form the replicase complex that bind to the sub-genomic promoter placed right before the S protein sequence. The cloned portions of all plasmid constructs were verified by DNA sequencing. Plasmids were linearized immediately after the poly(A) stretch and used as a template for *in vitro* transcription reaction with T7 RNA polymerase. For LUNAR-CoV19 vaccine, the reaction for RNA was performed as previously described [43] with proprietary modifications to allow highly efficient co-transcriptional incorporation of a proprietary Cap1 analogue and to achieve high quality RNA molecule of over 11,000-nt long the STARR mRNA. RNA was then purified through silica column (Macherey Nagel) and quantified by UV absorbance. For the conventional mRNA vaccine, the RNA was synthesized similarly but with 100% substitution of UTP with N1-methyl-pseudoUTP. For both LUNAR-CoV19 and conventional mRNA vaccines, the RNA quality and integrity were verified by 0.8-1.2% non-denaturing agarose gel electrophoresis as well as Fragment Analyzer (Advanced Analytical). The purified RNAs were stored in RNase-free water at −80 °C until further use.

### Vaccine lipid nanoparticles (LNPs)

LUNAR® nanoparticles encapsulating STARR™ were prepared by mixing an ethanolic solution of lipids with an aqueous solution of STARR™ RNA. Lipid excipients (Arcturus Therapeutics proprietary ionizable lipid, DSPC, Cholesterol and PEG2000-DMG) are dissolved in ethanol at mole ratio of 50:10: 38.5:1.5 or 50:13:35.5:1.5. An aqueous solution of the vaccine RNA is prepared in citrate buffer pH 4.0. The lipid mixture is then combined with the vaccine RNA solution at a flow rate ratio of 1:3 (V/V) via a proprietary mixing module. Nanoparticles thus formed are stabilized by dilution with phosphate buffer followed by HEPES buffer, pH 8.0. Ultrafiltration and diafiltration (UF/DF) of the nanoparticle formulation is then performed by tangential flow filtration (TFF) using modified PES hollow-fiber membranes (100 kDa MWCO) and HEPES pH 8.0 buffer. Post UF/DF, the formulation is filtered through a 0.2 μm PES filter. An in-process RNA concentration analysis is then performed. Concentration of the formulation is adjusted to the final target RNA concentration followed by filtration through a 0.2 μm PES sterilizing-grade filter. Post sterile filtration, bulk formulation is aseptically filled into glass vials, stoppered, capped, and frozen at −70 ± 10°C. Analytical characterization included measurement of particle size and polydispersity using dynamic light scattering (ZEN3600, Malvern Instruments), pH, Osmolality, RNA content and encapsulation efficiency by a fluorometric assay using Ribogreen RNA reagent, RNA purity by capillary electrophoresis using fragment analyzer (Advanced Analytical), lipid content using HPLC,.

### In vitro transfection and immunoblot detection of spike protein

Hep3b cells (seeded in 6-well plates at a density of 7 × 10^5^ cells/well, a day before) were transfected with purified IVTs (2.5 μg conventional mRNA and 2.5 μg STARR) by Lipofectamine MessengerMax transfection reagent (Thermo Fisher Scientific) according to the manufacturer’s instruction. The cells were harvested the next day with a hypotonic buffer (10 mM Tris-HCl, 10 mM NaCl supplemented with protease inhibitor cocktail (Roche)) followed by sonication. Samples were deglycosylated followed by treatment with PNGase F (New England Biolabs) according to the manufacture’s instruction.

The protein lysate (10 μg) was resolved on a 7.5% NuPAGE Tris-Acetate gel (Thermo Fisher Scientific), and the spike protein expression was analyzed by LI-COR Quantitative Western Blot system using a rabbit antibody detecting S1 (40150-T62-COV2, Sino Biologic) and a mouse antibody for S2 region (GTX632604, GeneTex) along with appropriate secondary antibodies (goat anti-rabbit 800 and goat anti-mouse 680).

### Animal studies

#### BALB/c studies

All BALB/c mouse studies were approved by the Explora Biolabs IACUC and performed under the Animal Care and Use Protocol number EB-17-004-003. A head-to-head comparison of the protein expression of the conventional mRNA and STARR vaccine platforms was conducted using conventional mRNA and STARR constructs expressing a luciferase reporter gene. BALB/c mice (Jackson Laboratory) were intramuscularly (IM) in the *rectus femoris* with conventional mRNA or STARR at doses of 0.2, 2 and 10 μg (*n*=3 mice/group). Expression of the conventional mRNA and STARR constructs were measured at days 1, 3 and 7 post-inoculation through luciferase expression by imaging the mice for bioluminescence.

Humoral responses to the SARS-CoV-2 Spike vaccine candidates were tested in Female BALB/c mice (Jackson Laboratory) aged 8-10 weeks by IM immunization of the *rectus femoris* with either conventional mRNA or LUNAR-COV19 at doses 0.2 μg, 2 μg, or 10 μg (*n*=5 mice/group). Mice were bled at baseline and at 10, 19, 30, 40, 50- and 60-days post-vaccination to assess SARS-CoV-2 specific humoral immune responses.

#### C57BL/6

All C57BL/6 mouse studies were performed in accordance with protocols approved by the Institutional Animal Care and Use Committee at Singapore Health Services, Singapore (ref no.: 2020/SHS/1554). C57BL/6 mice purchased from inVivos were housed in a BSL-2 animal facility at Duke-NUS Medical School. Groups of 6-8 weeks old wild-type C57BL/6 female mice were vaccinated intramuscularly with either conventional mRNA or LUNAR-COV19 at doses 0.2 μg, 2 μg, or 10 μg. For transcriptomic and T cell studies, submandibular bleeds were performed for whole blood at 24 hrs post-vaccination. Day 7 post-immunization, mice were sacrificed at and inguinal lymph nodes and spleen harvested for investigation of immune gene expression and T cell responses, respectively. Splenocyte suspensions for measuring T cell responses were obtained by crushing spleen through a 70μm cell strainer (Corning). Red blood cells were removed by lysis using BD PharmLyse reagent. For antibody studies, another set of vaccinated 6-8 weeks old mice were bled at baseline and at 10, 20, and 30 days post-vaccination.

SARS-CoV-2 challenge experiments were conducted with female B6;SJL-Tg(K18-hACE2)2Prlmn/J mice purchased from Jackson laboratory. Groups of 6-8 weeks old wild-type C57BL/6 female mice were vaccinated intramuscularly with 100 μl LUNAR-COV19 at doses of 2 μg, or 10 μg. Submandibular bleeds were performed for serum isolation to determine antibody titers via PRNT 28 days post vaccination. Animal were infected with 5×10^4^ TCID50 in 50μl via the intranasal route. Daily weight measurements and clinical scores were obtained. Mice were sacrificed when exhibiting greater than 20% weight loss or clinical score of 10. To assess organ viral loads, mice were sacrificed 5 days post infection and harvested organs were frozen at −80°C. Whole lungs and brains were homogenized with MP lysing matrix A and F according to manufacturer’s instructions in 1ml PBS. Homogenate was used to assess both plaque titers and RNA extraction using TRIzol LS (Invitrogen). No blinding was done for animal studies.

### Gene expression of immune and inflammatory genes

Whole blood collected 1-day post-vaccination was lysed using BD PharmLyse reagent and RNA extracted using Qiagen RNAeasy kit. Mouse lymph nodes collected from 7 days post vaccination were homogenized and RNA extracted using trizol LS. RNA (50 ng) from whole blood cells and lymph nodes were hybridized to the NanoString nCounter mouse inflammation and immunology v2 panels (Nanostring Technologies), respectively. As previously described [20, 44], RNA was hybridized with reconstituted CodeSet and ProbeSet. Reactions were incubated for 24 hours at 65°C and ramped down to down to 4°C. Hybridized samples were then immobilized onto a nCounter cartridge and imaged on a nCounter SPRINT (NanoString Technologies). Data was analyzed using the nSolver Analysis software (Nanostring Technologies) and Partek Genomics Suite. For normalization, samples were excluded when percentage field of vision registration is <75, binding density outside the range 0.1–1.8, positive control *R*2 value is <0.95 and 0.5 fM positive control is ≤2 s.d. above the mean of the negative controls. Background subtraction was performed by subtracting estimated background from the geometric means of the raw counts of negative control probes. Probe counts less than the background was floored to a value of 1. The geometric mean of positive controls was used to compute positive control normalization parameters. Samples with normalization factors outside 0.3–3.0 were excluded. The geometric mean of housekeeping genes was used to compute the reference normalization factor. Samples with reference factors outside the 0.10–10.0 range were also excluded. Hierarchical clustering was performed with Partek Genomics Suite v6 on gene sets zScore values by Euclidean dissimilarity and average linkage.

To identify DEGs between groups, Partek Genomics Suite Analysis v7 software was used to analyse variance (ANOVA) with a cut off-of *P* < 0.05. Log_2_ Fold Changes generated were used for volcano plots constructed using Prism v8.1.0 software. DEGs were identified by a fold change cut-off of 2. Unsupervised principle component analysis was performed to visualize variability between vaccinated and non-vaccinated animals with Partek genomics suite analysis v7 software. PCA ellipsoids were drawn with a maximum density and 3 subdivisions.

### Flow cytometry

Surface staining was performed on freshly-isolated splenocytes using the following panel of antibodies and reagents: B220 (RA3-6B2), CD3 (17A2), CD4 (RM4-5), CD8α (53-6.7), CD44 (IM7), CD62L (MEL-14) and DAPI. Intracellular cytokine staining was performed by stimulating freshly-isolated splenocytes with 50 ng/ml PMA and 500 ng/ml ionomycin in the presence of GolgiPlug (BD) for 6 h. After stimulation, surface staining of CD3, CD4 and CD8a was performed followed by intracellular staining of IFN-γ (XMG1.2) and IL-4 (11B11). Data acquisition was performed on a BD LSRFortessa and analyzed using FlowJo.

### ELISPOT

ELISPOT was performed using mouse IFN-γ ELISpot^BASIC^ kit (Mabtech). A similar protocol has been used for human SARS-CoV-2 samples [40]. In brief, 4 × 10^5^ freshly-isolated splenocytes were plated into PVDF-coated 96 well plates containing IFN-γ capture antibody (AN18). Cells were stimulated with a 15-mer peptide library covering part of the S protein. 143 total peptides were divided into four pools and used at a final concentration of 1 μg/ml per peptide. Negative control wells contained no peptide. Following overnight stimulation, plates were washed and sequentially incubated with biotinylated IFN-γ detection antibody (R4-6A2), streptavidin-ALP and finally BCIP/NBT. Plates were imaged using ImmunoSpot analyzer and quantified using ImmunoSpot software.

### Luminex Immuno-assay

#### Longitudinal assessment of binding antibody

Longitudinal IgM and IgG responses in BALB/c and C57BL/6 were measured using an in-house Luminex Immuno-assay. Similar Luminex Immuno-assays have been previously described for antibody detection against SARS-CoV-2 antigens [45, 46]. Briefly, Magpix Luminex beads were covalently conjugated to insect-derived HIS-tagged SARS-CoV-2 whole Spike protein (SinoBiologicals) using the ABC coupling kit (Thermo) as per manufacturer’s instructions. Beads were then blocked with 1%BSA, followed by incubation with serum (diluted at 1:2000 in block) for 1 hr at 37C. Beads are then washed and SARS-CoV-2 Spike-specific mouse antibodies were detected using the relevant biotinylated secondary antibody (i.e. anti-mouse IgM-biotin and anti-mouse IgG-biotin (Southern Biotech) for IgM and IgG assessment, respectively) with streptavidin-PE (Southern Biotech). Antibody binding to Spike were then measured on a Magpix instrument as median fluorescence intensity (MFI). Spike antigen quantity on beads were also probed with anti-6xHIS-PE antibodies and all MFI values were then corrected to Spike antigen quantity to account for experiment to experiment variation.

#### IgG and IgG subclass endpoint titers

IgG endpoint titers to mammalian-derived SARS-CoV-2 Spike, Spike domain 1 (S1), spike domain 2 (S2) and receptor binding domain (RBD) at day 30 sera post-immunization were measured using Luminex ImmunoAssay. Assay was conducted as described above, with the modification of serially diluting serum 10-fold from 200 to 2×10^8^. Similarly, IgG subclass endpoint titers (i.e. IgG1 and IgG2a in BALB/c and IgG1 and IgG2c in C57BL/6) were measured against mammalian-derived SARS-CoV-2 Spike protein using serially diluted mouse sera (5-fold from 200 to 3.1×10^6^) and secondary antibodies anti-IgG1-biotin, anti-IgG2a-biotin or anti-IgG2b-biotin (Southern Biotech). Four parameter logistic (4PL) curves were fitted to the measured MFI data from serially diluted sera, and three times the background (i.e. 3x MFI with no serum) was used as a threshold cutoff to estimate endpoint titers.

#### IgG Avidity

Avidity index of IgG to SARS-CoV-2 Spike protein at day 30 sera post-immunization was estimated using the Luminex ImmunoAssay. Assay was conducted as described above with the minor modification of following bead incubation with serum (diluted at 1:2000) with either a 10 min PBS or 8M urea wash. Avidity Index was estimated by subtracting background MFI from all sample values, and then dividing MFI with 8M Urea wash by MFI with PBS wash.

### Neutralization assay

#### Virus Neutralization titer assay (VNT)

Neutralization sero-conversion was assessed at day 10 and 20 post-immunization in BALB/c using a virus neutralization assay as previously described [47]. Briefly, sera were diluted to 1:20 in culture media, mixed at a 1:1 ratio with a Singaporean clinical isolate of live SARS-CoV-2 virus, isolate BetaCoV/Singapore/2/2020 (GISAID accession code EPI_ISL_406973) and incubated for 1 hr at 37C. Virus-antibody immune-complexes were then added to Vero-E6 cells (ATCC) in 96-well plates, and incubated at 37C. Five days later, plates were assessed under a microscope for cytopathic effect (CPE) of the cells.

#### Plaque reduction neutralization titer (PRNT)

Neutralization of live SARS-CoV-2 was measured by PRNT at day 30 post-vaccination in both BALB/c and C57BL/6 mice. Similar protocols have been published previously for SARS-CoV-2 [48]. Briefly, mouse sera were serially diluted from 1:20 to 1:320 in culture media and incubated with the Singapore isolate of SARS-CoV-2 virus for 1 hr at 37C. Virus-antibody mixtures were then added to Vero-E6 cells in 24-well plates, incubated for 1-2 hrs, then overlayed with carboxymethyl cellulose (CMC) and incubated at 37C under 5% CO_2_. At 5 days, cells are washed, stained with crystal violet and plaques counted. The serum dilution leading to neutralization of 50% of virus, i.e. PRNT_50_, was estimated.

## ACKNOWLEDGMENTS

We thank the Economic Development Board of Singapore for initiating this collaboration and for funding the development of LUNAR-COV19. ARdA received salary support from the National Medical Research Council (NMRC) Young Investigator Award. EEO received salary support from the NMRC Clinician-Scientist Award (Senior Investigator).

## AUTHOR CONTRIBUTIONS

Ruklanthi de Alwis was responsible for the humoral characterization of the immune response from the LUNAR COV19 vaccine. Esther S Gan was responsible for the huACE2 transgenic mouse challenge studies and expression profiling analysis of the LUNAR COV19 vaccine.

---

**Supplementary Figure 1.**
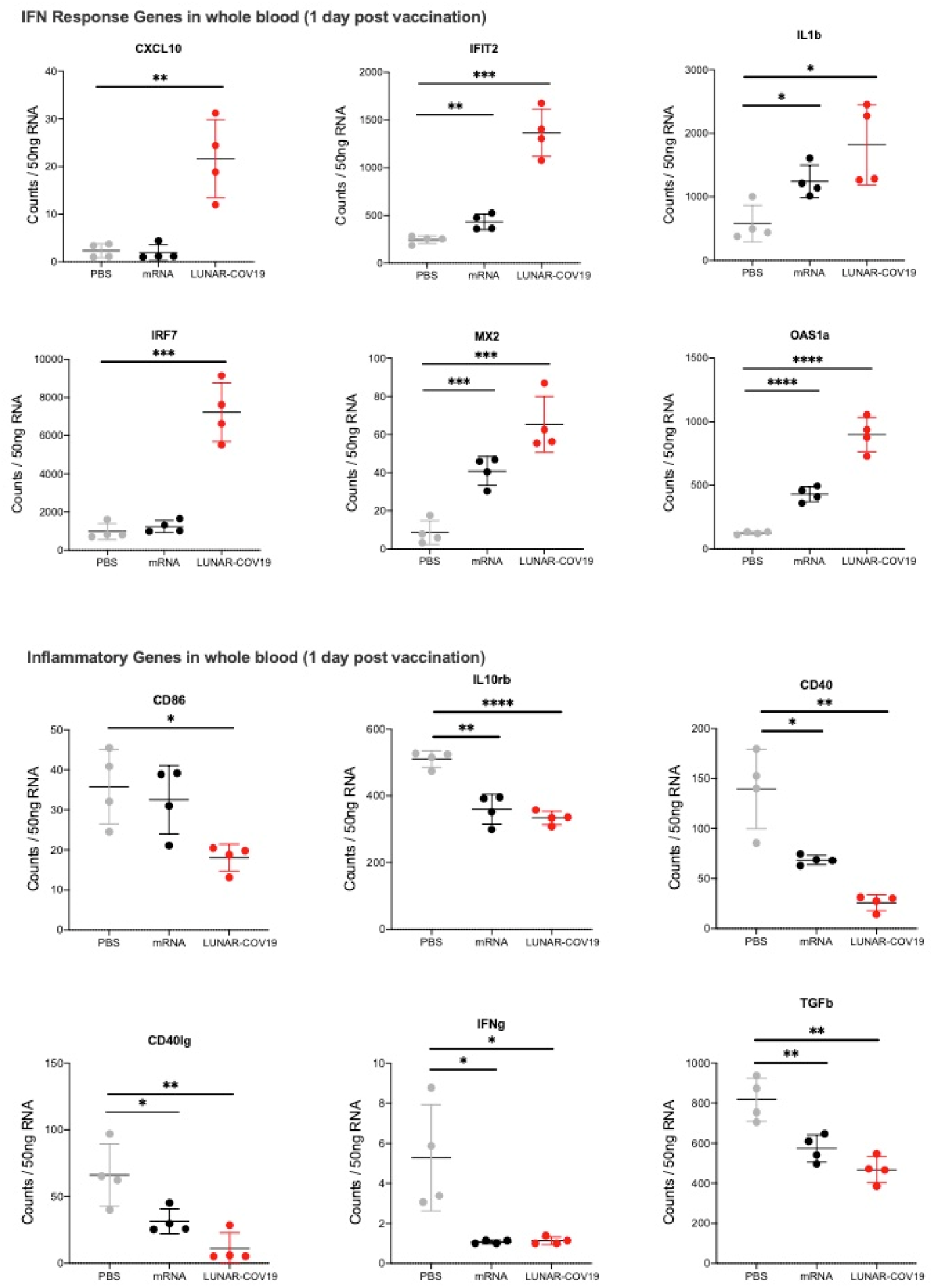
Whole blood transcriptomic data at 1-day post-prime vaccination showing Nanostring counts per 50ng RNA of selected IFN and inflammatory genes.

**Supplementary Figure 2.**
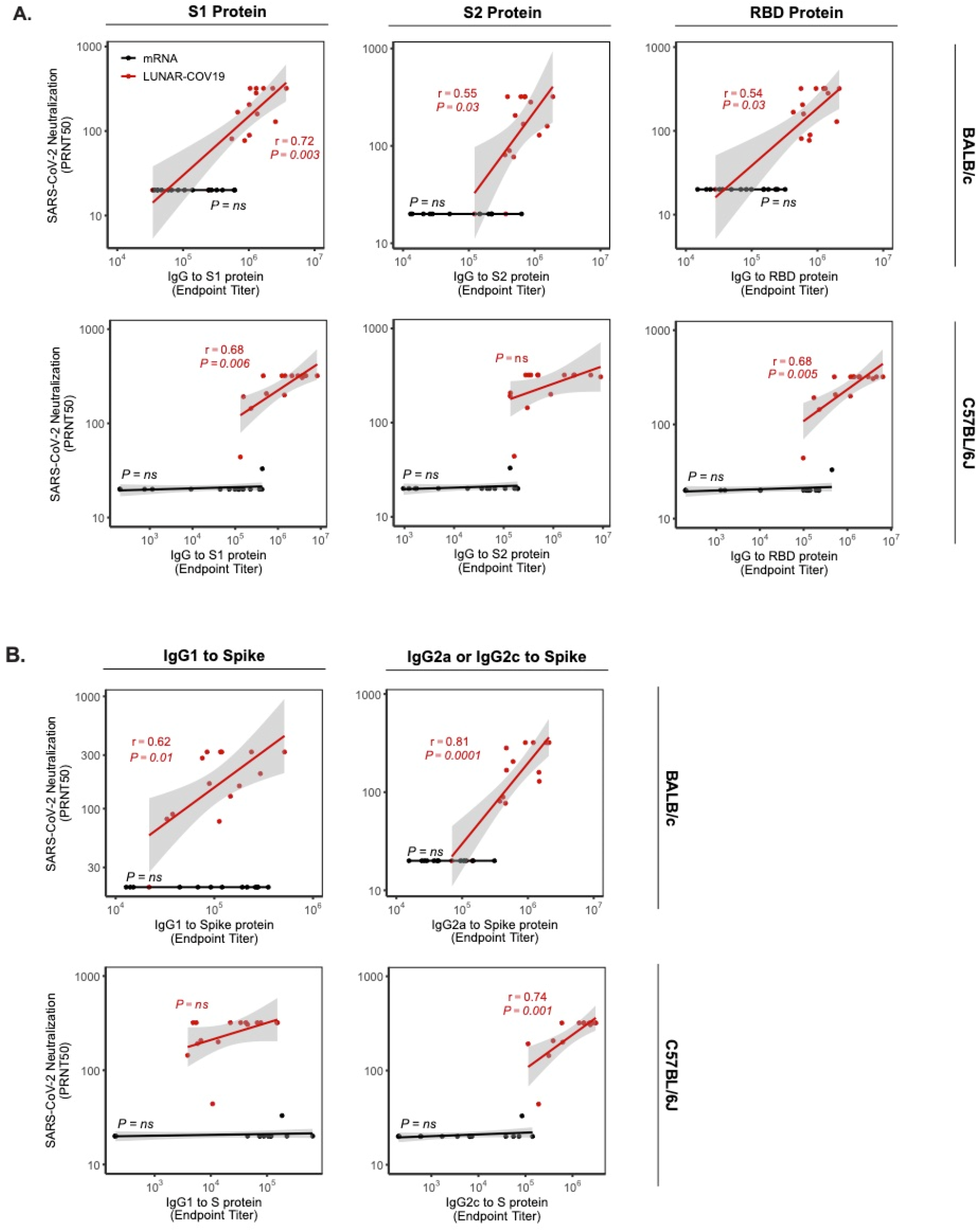
Correlation analysis of live SARS-CoV-2 neutralization against binding IgG and IgG subclasses in BALB/c and C57BL/6J mouse strains. **A**) Spearman correlation analysis of SARS-CoV-2 neutralization (PRNT50) against total IgG specific to several SARS-CoV2 antigens, including S, S1, and RBD recombinant proteins. **B**) Spearman correlation analysis of SARS-CoV-2 neutralization (PRNT50) against SARS-CoV2 S-specific IgG subclasses (IgG1 and IgG2a or IgG2c).

